# Phylogenomic species delimitation in microendemic frogs of the Brazilian Atlantic Forest

**DOI:** 10.1101/143735

**Authors:** Marcio R. Pie, Marcos R. Bornschein, Luiz F. Ribeiro, Brant C. Faircloth, John E. McCormack

**Affiliations:** Departamento de Zoologia, Universidade Federal do Paraná, CEP 81531–980, Curitiba, Paraná, Brazil; Mater Natura – Instituto de Estudos Ambientais, CEP 80250–020, Curitiba, Paraná, Brazil; Instituto de Biociências, Universidade Estadual Paulista, Praça Infante Dom Henrique s/no, Parque Bitaru, CEP 11330–900, São Vicente, São Paulo, Brazil; Escola de Ciências da Vida, Pontifícia Universidade Católica do Paraná, CEP 80215–901, Curitiba, Paraná, Brazil; Department of Biological Sciences and Museum of Natural Science, Louisiana State University, Baton Rouge, LA 70803, USA; Moore Laboratory of Zoology, Occidental College, 1600 Campus Road, Los Angeles, California 90041, USA

**Keywords:** Anura, Brazil, Terrarana, Atlantic Forest, *Brachycephalus*, *Melanophryniscus*

## Abstract

The advent of next-generation sequencing allows researchers to use large-scale datasets for species delimitation analyses, yet one can envision an inflection point where the added accuracy of including more loci does not offset the increased computational burden. One alternative to including all loci could be to prioritize the analysis of loci for which there is an expectation of high informativeness, such as those with higher numbers of parsimony-informative sites. Here, we explore the issue of species delimitation and locus selection with species from two anuran genera: *Melanophryniscus* (Bufonidae) and *Brachycephalus* (Brachycephalidae). Montane species in these genera have been isolated in sky islands across the southern Brazilian Atlantic Forest, which led to the formation of a number of microendemic species. To delimit species, we obtained genetic data using target enrichment of ultraconserved elements from 32 populations (13 for *Melanophryniscus* and 19 for *Brachycephalus*), and we were able to create datasets that included over 800 loci with no missing data. We ranked loci according to their corresponding number of parsimony-informative sites, and we performed species delimitation analyses using BPP in each genus based on the top 10, 20, 40, 80, 160, 320, and 640 loci. We also conducted several additional analyses using 10 randomly sampled datasets containing the same numbers of loci to discriminate the relative contribution of increasing the number of loci from prioritizing those with higher informativeness. We identified three types of node: nodes with either consistently high or low support regardless of the number of loci or their informativeness, and nodes that were initially poorly supported, but their support became stronger with more data. Adding more loci had a stronger impact on model support than prioritizing loci for their informativeness, but this effect was less apparent in datasets with more than 160 loci. When viewed across all sensitivity analyses, our results suggest that the current species richness in both genera might have been underestimated. In addition, our results provide useful guidelines to the use of different sampling strategies to carry out species delimitation with phylogenomic datasets.

---

Accurate species delimitation forms the basis of much of biodiversity research (Sites and Marshall 2004; Adams et al. 2014). Two important advances in this area have been obtained in recent years. The first is the development of species delimitation methods (e.g. Yang 2002; Rannala and Yang 2003; Knowles and Carstens 2007; Yang and Rannala 2010; Ence and Carstens 2011; see Rannala 2015 for a recent review) that are based on the multispecies coalescent (MSC) model (Takahata et al. 1995; Rannala and Yang 2003). These methods provide an objective and operational way to infer species limits that is explicitly based on a rigorous population genetic framework (Fujita and Leaché 2011; Rannala 2015; but see Sukumaran and Knowles 2017). The second involves advances in sequencing technologies that allow for the generation of large-scale datasets (Bi et al. 2012; Faircloth et al. 2012; Lemmon et al. 2012; Lemon and Lemon 2013; McCormack et al. 2013; Smith et al. 2013). These technologies allow researchers to collect data from hundreds or thousands of loci across hundreds or thousands of species in an efficient manner. Unfortunately, the computational demands of MSC species delimitation methods when dealing with these large datasets means that the brute-force approach of including as many loci as possible might not be the most computationally efficient and/or cost effective. The ideal approach might, instead, be to reduce the total number of loci included by focusing individual analyses on more informative loci while excluding those with low information content because this latter class of loci increases the computational burden while contributing little information to the analysis. In the analogous case of species tree inference under the MSC, some methods perform worse with the addition of low-information loci (e.g. Manthey et al. 2016; Xu and Yang 2016; but see Xi et al. 2015). Conversely, some recent speciation events might need a large number of loci to be properly detected, suggesting that researchers should use the largest number of loci possible. Central to this question is how varying the number and their informativeness affects performance of species delimitation methods. To the best of our knowledge, the only empirical study to assess locus number and informativeness on species delimitation was conducted by Hime et al. (2016) using populations of the stream-dwelling salamander *Ambystoma ordinarium* with datasets between 10 and 89 loci.

Here, we investigate the performance of MSC species delimitation when applied to loci collected from co-distributed montane species within two anuran genera: *Melanophryniscus* (Bufonidae) and *Brachycephalus* (Brachycephalidae). The genus *Melanophryniscus* is broadly distributed throughout southeastern South America, spanning south and southeastern Brazilian Atlantic Forest, wetlands and grasslands of Brazil, inter-Andean valleys in Bolivia, and areas across Paraguay and Uruguay down to central Argentina (Frost 2017). Importantly, *Melanophryniscus* of the southern Brazilian Atlantic Forests are characterized by montane endemic species with restricted and isolated distributions in cloud forests, *campos de altitude*, and grasslands (Langone et al. 2008; Steinbach-Padilha 2008; Bornschein et al. 2015). These include five of the 29 currently described *Melanophryniscus* species (Frost 2017): *M. alipioi, M. biancae, M. milanoi, M. vilavelhensis*, and *M. xanthostomus*. Of those species, *M. biancae* and *M. vilavelhensis* represent a distinct lineage within montane *Melanophryniscus*, given their phylogenetic distance from the remaining species (Firkowski et al. 2016) and the unique type of vegetation in which they are found (Bornschein et al. 2015). The remaining three species *(M. alipioi, M. milanoi*, and *M. xanthostomus*) could be species complexes (Firkowski et al. 2016), but the considerable morphological variability found among individuals of these species, even within a given location, is a major hurdle for the delimitation of potential species using phenotypic data alone. Members of the genus *Brachycephalus* are endemic to the Brazilian Atlantic Forest, with a distribution extending nearly 1700 km along the biome, and most *Brachycephalus* species occur in isolated mountaintops from the Brazilian states of Bahia in northeastern Brazil to Santa Catarina in southern Brazil (Pie et al. 2013; Bornschein et al. 2016a). *Brachycephalus* includes both cryptic and aposematic toadlets, which live in the forest leaf litter and are commonly active during the day (e.g. Ribeiro et al. 2015). The most obvious feature of this genus is their extreme level of miniaturization (SVL≈1-1.5 cm), which led to extreme modifications to their life histories (Hanken and Wake 1993; Yeh 2002). The genus was recently divided into three species groups (namely the *pernix, ephippium*, and *didactylus* groups - see Ribeiro et al. [2015]), with the *pernix* group currently including 15 described species distributed across the southern Atlantic Forest (Bornschein et al. 2016a). Species of *Brachycephalus* and *Melanophryniscus* have traditionally been described using phenotypic data alone (Bornschein et al. 2015, 2016b; Pie and Ribeiro 2015; Ribeiro et al. 2015), which can lead to underestimates of species diversity (Bickford et al. 2007). A recent delimitation study using a handful of loci suggested that there could be a number of *Brachycephalus* and *Melanophryniscus* species that remain undescribed, although variability between loci in delimitation schemes (Firkowski et al. 2016) suggested that more extensive data were needed to firmly establish species boundaries.

The main goal of this study is to delimit species of *Brachycephalus* and *Melanophryniscus* under the MSC using ultraconserved elements (UCEs, Faircloth et al. 2012). Specifically, we used a Bayesian Markov chain Monte Carlo program for species delimitation (BPP 3.3; Yang 2015) that relies on the MSC model to compare different models of species delimitation while accounting for incomplete lineage sorting due to ancestral polymorphism and gene tree-species tree conflicts (Yang and Rannala 2010, 2014; Rannala and Yang 2013). The lineages investigated in this study are particularly suitable for the use of BPP because the candidate species are almost always allopatric, such that gene flow among populations is likely low to nonexistent (simulation studies suggest that the migration threshold for genetic isolation is about 1 individual per 10 generations [Zhang et al. 2011]). In addition to investigating species limits in *Brachycephalus* and *Melanophryniscus*, we also explored the importance of locus variation (informativeness) and locus number on the ideal species delimitation schemes.

## Methods

We obtained tissue samples from field-collected specimens of 13 populations of *Melanophryniscus* and 19 populations of *Brachycephalus* (Table 1), and we deposited voucher specimens in the herpetological collection of the Department of Zoology of the Universidade Federal do Paraná (DZUP) in Curitiba, Brazil (more information on specimen collection methods and localities can be found in Firkowski et al. [2016]). These samples include most described species of *Brachycephalus* of the *pernix* group, except for *B. albolineatus*, *B. leopardus*, and *B. tridactylus*. *Brachycephalus sulfuratus* does not belong to the *pernix* species group, but was included in the analyses to improve the rooting of the guide tree. Also, these samples include all known records of montane *Melanophryniscus*, except those ascribed to *M. biancae* and *M. vilavelhensis*, including several novel records in relation to Bornschein et al. (2015). We extracted genomic DNA using PureLink Genomic DNA kit (Invitrogen, USA) and we fragmented the obtained DNA using a BioRuptor NGS (Diagenode). We prepared Illumina libraries using KAPA library preparation kits (Kapa Biosystems) and custom sequence tags unique to each sample (Faircloth and Glenn 2012). To enrich targeted UCE loci, we followed an established workflow (Gnirke et al. 2009; Blumenstiel et al. 2010) while incorporating several modifications to the protocol detailed in Faircloth et al. (2012). Specifically, we pooled eight samples at equimolar ratios, prior to enrichment, and we blocked the Illumina TruSeq adapter sequence using custom blocking oligos. We enriched each pool using a set of 2,560 custom-designed probes (MYcroarray, Inc.) targeting 2,386 UCE loci (see Faircloth et al. [2012] and http://ultraconserved.org [last accessed May 23, 2017] for details on probe design). Prior to sequencing, we qPCR-quantified enriched pools, combined pools at equimolar ratios, and sequenced the combined libraries using two partial (50%) runs of a MiSeq PE250 (Cofactor Genomics).

**Table 1.**
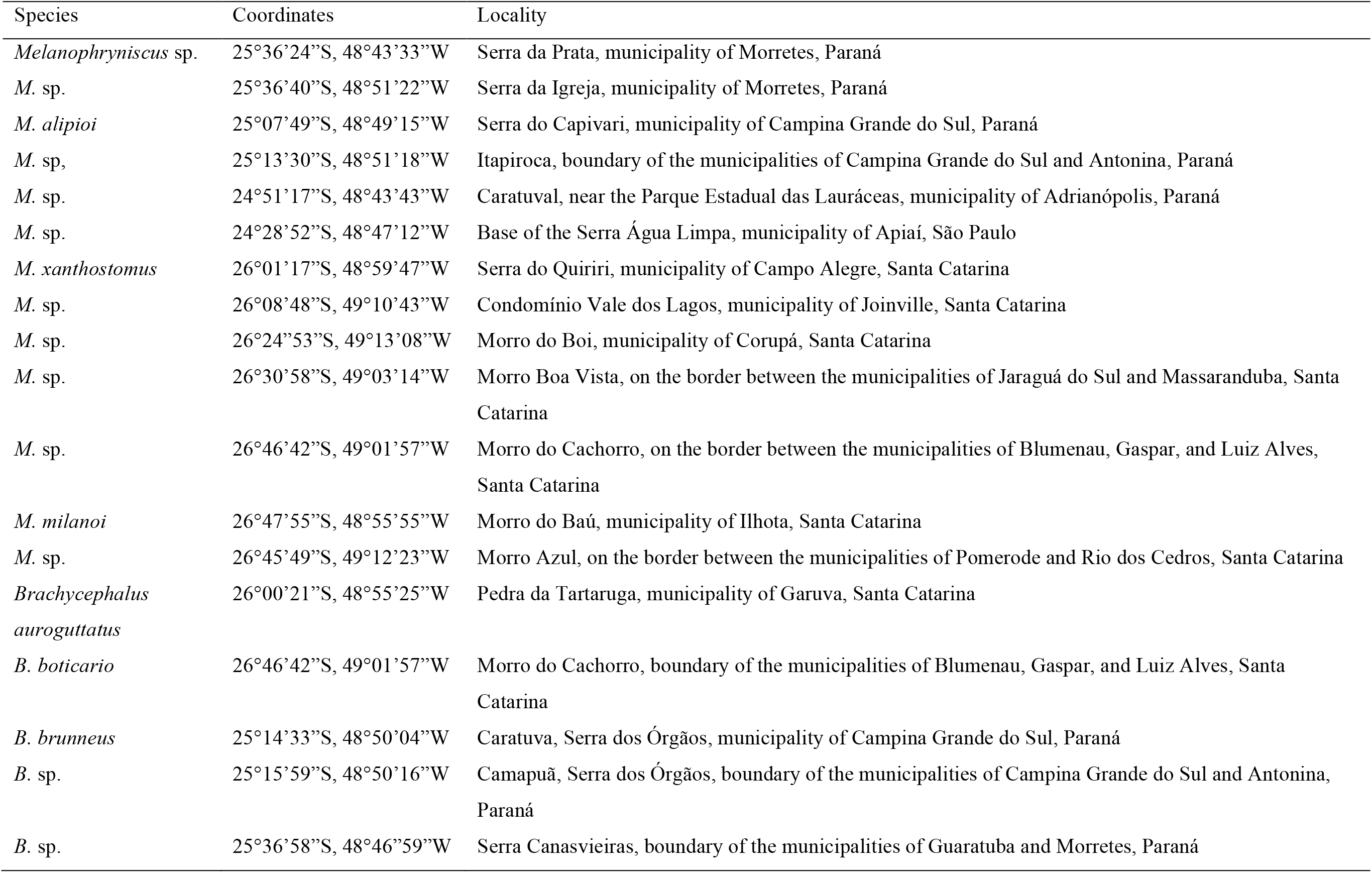
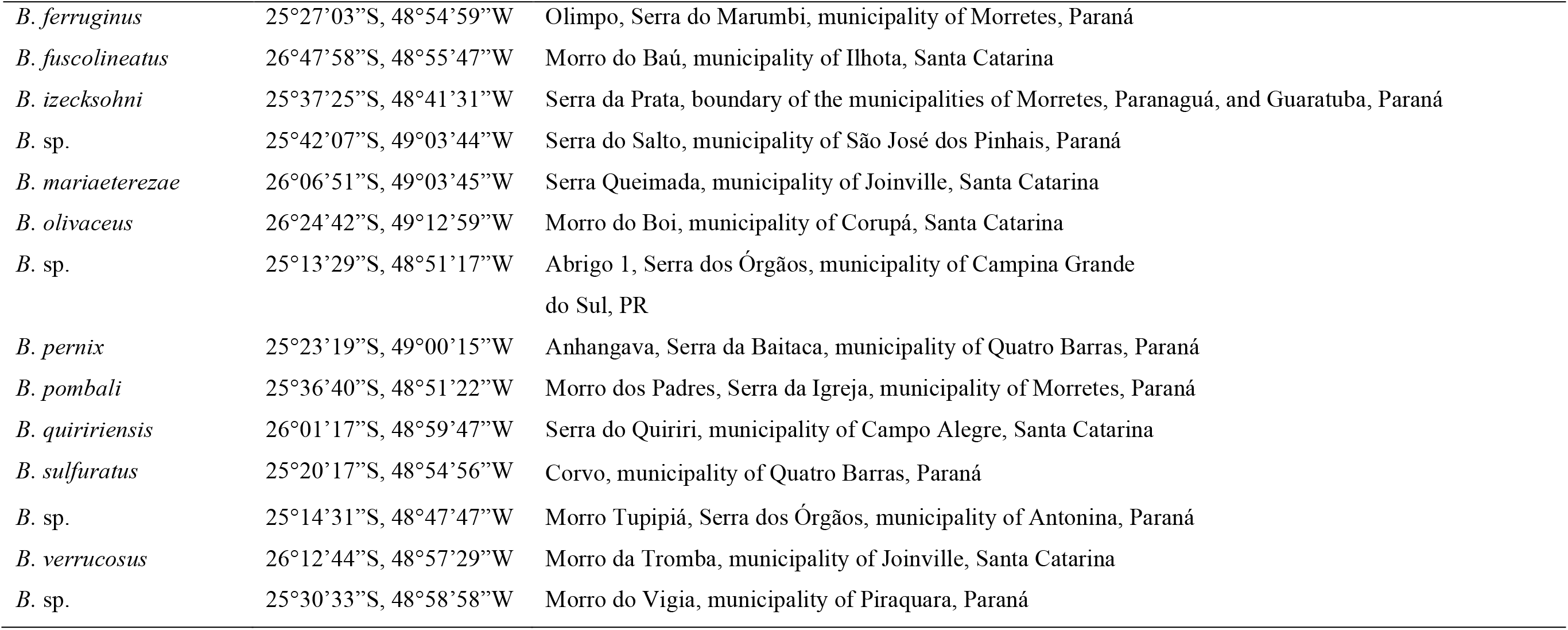
Samples used in the present study, with the corresponding localities and geographical coordinates.

| Species | Coordinates | Locality |
| --- | --- | --- |
| *Melanophryniscus* sp. | 25°36'24"S, 48°43'33"W | Serra da Prata, municipality of Morretes, Paraná |
| *M.* sp. | 25°36'40"S, 48°51'22"W | Serra da Igreja, municipality of Morretes, Paraná |
| *M. alipioi* | 25°07'49"S, 48°49'15"W | Serra do Capivari, municipality of Campina Grande do Sul, Paraná |
| *M.* sp. | 25°13'30"S, 48°51'18"W | Itapiroca, boundary of the municipalities of Campina Grande do Sul and Antonina, Paraná |
| *M.* sp. | 24°51'17"S, 48°43'43"W | Caratuval, near the Parque Estadual das Lauráceas, municipality of Adrianópolis, Paraná |
| *M.* sp. | 24°28'52"S, 48°47'12"W | Base of the Serra Água Limpa, municipality of Apiaí, São Paulo |
| *M. xanthostomus* | 26°01'17"S, 48°59'47"W | Serra do Quiriri, municipality of Campo Alegre, Santa Catarina |
| *M.* sp. | 26°08'48"S, 49°10'43"W | Condomínio Vale dos Lagos, municipality of Joinville, Santa Catarina |
| *M.* sp. | 26°24'53"S, 49°13'08"W | Morro do Boi, municipality of Corupá, Santa Catarina |
| *M.* sp. | 26°30'58"S, 49°03'14"W | Morro Boa Vista, on the border between the municipalities of Jaraguá do Sul and Massaranduba, Santa Catarina |
| *M.* sp. | 26°46'42"S, 49°01'57"W | Morro do Cachorro, on the border between the municipalities of Blumenau, Gaspar, and Luiz Alves, Santa Catarina |
| *M. milanoi* | 26°47'55"S, 48°55'55"W | Morro do Baú, municipality of Ilhota, Santa Catarina |
| *M.* sp. | 26°45'49"S, 49°12'23"W | Morro Azul, on the border between the municipalities of Pomerode and Rio dos Cedros, Santa Catarina |
| *Brachycephalus auroguttatus* | 26°00'21"S, 48°55'25"W | Pedra da Tartaruga, municipality of Garuva, Santa Catarina |
| *B. boticario* | 26°46'42"S, 49°01'57"W | Morro do Cachorro, boundary of the municipalities of Blumenau, Gaspar, and Luiz Alves, Santa Catarina |
| *B. brunneus* | 25°14'33"S, 48°50'04"W | Caratuva, Serra dos Órgãos, municipality of Campina Grande do Sul, Paraná |
| *B.* sp. | 25°15'59"S, 48°50'16"W | Camapuã, Serra dos Órgãos, boundary of the municipalities of Campina Grande do Sul and Antonina, Paraná |
| *B.* sp. | 25°36'58"S, 48°46'59"W | Serra Canasvieiras, boundary of the municipalities of Guaratuba and Morretes, Paraná |
| *B. ferruginus* | 25°27'03"S, 48°54'59"W | Olimpo, Serra do Marumbi, municipality of Morretes, Paraná |
| *B. fuscolineatus* | 26°47'58"S, 48°55'47"W | Morro do Baú, municipality of Ilhota, Santa Catarina |
| *B. izecksohni* | 25°37'25"S, 48°41'31"W | Serra da Prata, boundary of the municipalities of Morretes, Paranaguá, and Guaratuba, Paraná |
| *B. sp.* | 25°42'07"S, 49°03'44"W | Serra do Salto, municipality of São José dos Pinhais, Paraná |
| *B. mariaeterezae* | 26°06'51"S, 49°03'45"W | Serra Queimada, municipality of Joinville, Santa Catarina |
| *B. olivaceus* | 26°24'42"S, 49°12'59"W | Morro do Boi, municipality of Corupá, Santa Catarina |
| *B. sp.* | 25°13'29"S, 48°51'17"W | Abrigo 1, Serra dos Órgãos, municipality of Campina Grande do Sul, PR |
| *B. pernix* | 25°23'19"S, 49°00'15"W | Anhangava, Serra da Baitaca, municipality of Quatro Barras, Paraná |
| *B. pombali* | 25°36'40"S, 48°51'22"W | Morro dos Padres, Serra da Igreja, municipality of Morretes, Paraná |
| *B. quiririensis* | 26°01'17"S, 48°59'47"W | Serra do Quiriri, municipality of Campo Alegre, Santa Catarina |
| *B. sulfuratus* | 25°20'17"S, 48°54'56"W | Corvo, municipality of Quatro Barras, Paraná |
| *B. sp.* | 25°14'31"S, 48°47'47"W | Morro Tupipiá, Serra dos Órgãos, municipality of Antonina, Paraná |
| *B. verrucosus* | 26°12'44"S, 48°57'29"W | Morro da Tromba, municipality of Joinville, Santa Catarina |
| *B. sp.* | 25°30'33"S, 48°58'58"W | Morro do Vigia, municipality of Piraquara, Paraná |

We filtered reads for adapter contamination, low-quality ends, and ambiguous bases using an automated pipeline (https://github.com/faircloth-lab/illumiprocessor) that incorporates Trimmomatic (Bolger et al. 2014). We assembled reads for each taxon using Trinity (Grabherr et al. 2011). We used the PHYLUCE software package (Faircloth 2015) to align assembled contigs back to their associated UCE loci, remove duplicate matches, and create a taxon-specific database of contig-to-UCE matches. We then generated two alignments: all *Brachycephalus* samples using *Melanophryniscus alipioi* as their outgroup, and all *Melanophryniscus* species using *Brachycephalus sulfuratus* as their outgroup. We selected loci to create 100% complete data sets for both genera, leading to 820 loci in the *Brachycephalus* data set and 1227 loci in the *Melanophryniscus* data set. We aligned data for each individual in each data set using MAFFT (Katoh 2013), and we trimmed resulting alignments using GBlocks (Castresana 2000) with default parameters.

Ideally one would determine the informativeness of a given locus based on how much information it contributes to a given analysis (e.g. Townsend 2007; Gilbert et al. 2015). However, that would involve first running the entire analysis to then determine their informativeness *a posteriori*. A simple alternative is to calculate the absolute number of parsimony-informative sites (PIS – a single nucleotide polymorphism that is present in more than one individual) for each locus and to use this measure as a proxy for informativeness during species delimitation analyses, as recently proposed by Hime et al. (2016). Other studies used PISs as a measure of informativeness in the context of phylogenetic inference as well (e.g. Hosner et al. 2016; Meiklejohn et al. 2016). We calculated the number of PIS for each locus using ips 0.0-7 (Heibl 2014) and compared them between datasets using a Spearman’s test with the cor.test function in R 3.3.2 (R Core Team 2017). We then ranked loci in each dataset according to their corresponding number of PIS, and we created subsets of the top 10, 20, 40, 80, 160, 320, and 640 most informative loci for subsequent analyses.

We used BPP 3.3 (Yang 2015) in all species delimitation analyses. Important assumptions of the MSC model implemented in BPP are no recombination within a locus, free recombination between loci, no migration (gene flow) between species and neutral evolution, and all of which seem reasonable given our model system of allopatric populations assessed using UCEs. It is also important to note that, although the core of UCEs tend to be highly conserved due to strong selection, flanking regions tend to behave similarly to neutral DNA from introns (see Jarvis et al. 2014). Every population was considered initially as a potential species as a conservative starting point (see Olave et al. 2014), allowing for the possibility of later being lumped together by BPP depending on the analyzed data. Three different sets of priors for population size (θ) and divergence time at the root of the species tree (τ0) being assigned gamma priors were used: (1) small ancestral population sizes and shallow interspecific divergences: θ ~ Γ(2, 1000), τ_0_~ Γ(2, 1000); (2) large ancestral population sizes and shallow interspecific divergences: θ ~ Γ(2, 1000), τ_0_~ Γ(1, 10); and (3) large ancestral population sizes and deep interspecific divergences: θ ~ Γ(1, 10), τ_0_~ Γ(1, 10). These sets of priors were used to test their effect on the chosen species delimitation scheme, but given what is known about the microendemic distribution and recent divergence times of the studied species, the first set of priors seems more realistic in this case and will be given priority when conflicting results are detected. Other divergence time parameters are assigned the Dirichlet prior (Yang and Rannala 2010: equation 2). We used the A10 model (speciesdelimitation=1, speciestree=0) for species delimitation using a user-specified guide tree (Yang and Rannala 2010; Rannala and Yang 2013). For computational reasons, we assumed that all loci have the same mutation rate (“locusrate=0”). We used the uniform rooted tree prior (speciesmodelprior=1) and the same θs were assumed across loci (heredity=0). We used the option of automatic fine-tuning the MCMC searches, which provided adequate acceptance proportions (i.e. between 20 and 80%). Analyses were repeated using rjMCMC algorithm 0 (ε=2) and algorithm 1 (α=2, m=1), as well as including/excluding sites with gaps (cleandata=0,1), given that the latter has been recently shown as potentially affecting species delimitation using phylogenomic data (Domingos et al. 2017). Each analysis was run at least twice to confirm consistency between runs, given that it seems to be the most effective method to guarantee the reliability of the results (Yang 2015). Given the number of tips on the guide tree, in each analysis there were 768 potential species delimitation models for *Brachycephalus* and 145 for *Melanophryniscus*. In total, we ran 336 analyses (2 genera X 7 datasets with varying numbers of loci X 2 algorithms X 3 sets of priors X 2 replicates X 2 treatments of gaps). Each analysis was run for 10,000 generations, sampling every 10^th^ generation and with the first 10% used as burnin. Although we are aware that BPP can carry out simultaneous species tree and species delimitation analyses (i.e. the A11 model), preliminary analyses using a variety of phylogenetic inference methods showed consistent topologies, such that the computational demands of coestimating species trees (e.g. Caviedes-Solis et al. 2015) would not compensate for the substantial increase in run time. Moreover, BPP has been shown to be robust to errors in guide trees (Zhang et al. 2014; Caviedes-Solis et al. 2015).

Finally, in order to discriminate the relative contribution of increasing the number of loci from prioritizing those with higher informativeness, 10 randomly-sampled datasets with the same numbers of loci were analyzed under model A10, algorithm 0, cleandata=0, and priors θ ~ Γ(2, 1000), τ_0_~ Γ(2, 1000), and the obtained results were contrasted with the corresponding previous analysis using ranked loci.

## Results

We found considerable variation among UCE loci in their informativeness, as indicated by the distribution of the number of PISs for each locus (Figure S1). In particular, most loci have few PIS and thus are relatively less able to provide information about intra- and interspecific genetic variation. Interestingly, when the same UCE locus was compared between the two datasets, we found a significant correlation in the number of PISs (ρ=0.31, p=1.88e-15), meaning that an informative locus from one genus tended to be informative in the other genus as well. This relationship was skewed, such that a poorly informative locus from one genus was usually also poorly informative in the other genus; however, a highly informative locus from one genus might not be equally informative in the other (Figure 1). Nevertheless, these results suggest that one could, in principle, select UCE loci that are most informative based on similar datasets involving lineages that are not very closely related (e.g. within Anura).

**Figure 1.**
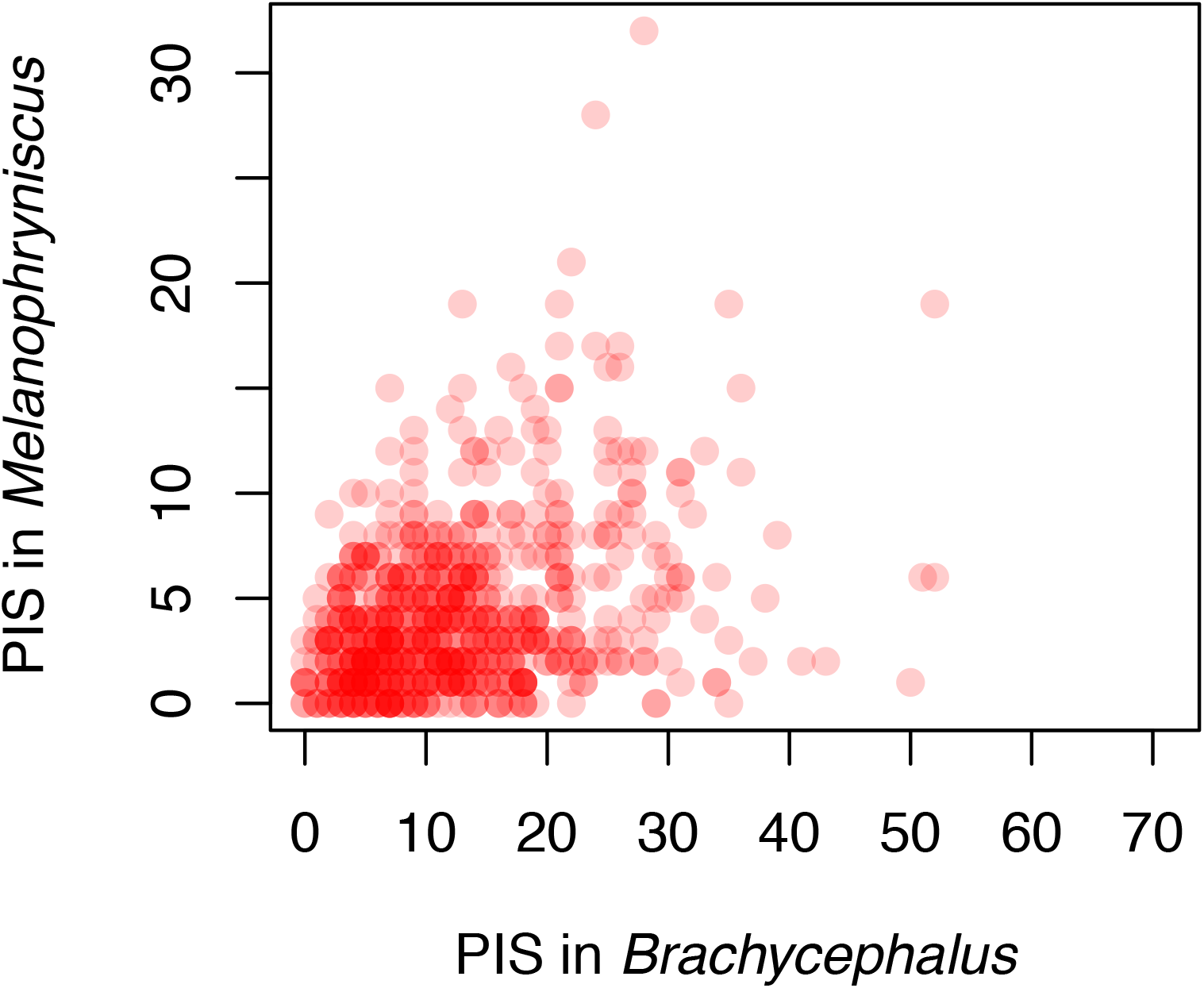
Correspondence between the number of parsimony-informative sites (PISs) for the same loci in the *Brachycephalus* and *Melanophryniscus* datasets.

A summary of the species delimitation analyses for *Melanophryniscus* and *Brachycephalus* are shown in Figures 2 and 3. There were three main types of support for splits among populations: (1) nodes showing 1.0 posterior probabilities (PPs) in all analyses, which were usually those closest to the base of the guide trees (2) nodes with low (or at least highly inconsistent) support across analyses, which were more often found in some regions near the tips of the guide trees; finally, (3) nodes ranging from low to very high support, which are indicated in Figures 2 and 3 with black circles. In the latter case, a clear pattern emerges of a monotonic increase in support with more loci (Figure 4). There seems to be a general monotonic increase in support with more data, which is expected in statistically consistent methods, although there was a slight decrease in the posterior probabilities between 20 and 40 for some analyses (e.g. Figure 4 E, F, L). Also, excluding sites with gaps (clear=1) does not seem to provide important insight, given that they simply led to more unstable posterior probability estimates (possibly because problematic sites had already been removed using Gblocks prior to the species delimitation).

**Figure 2.**
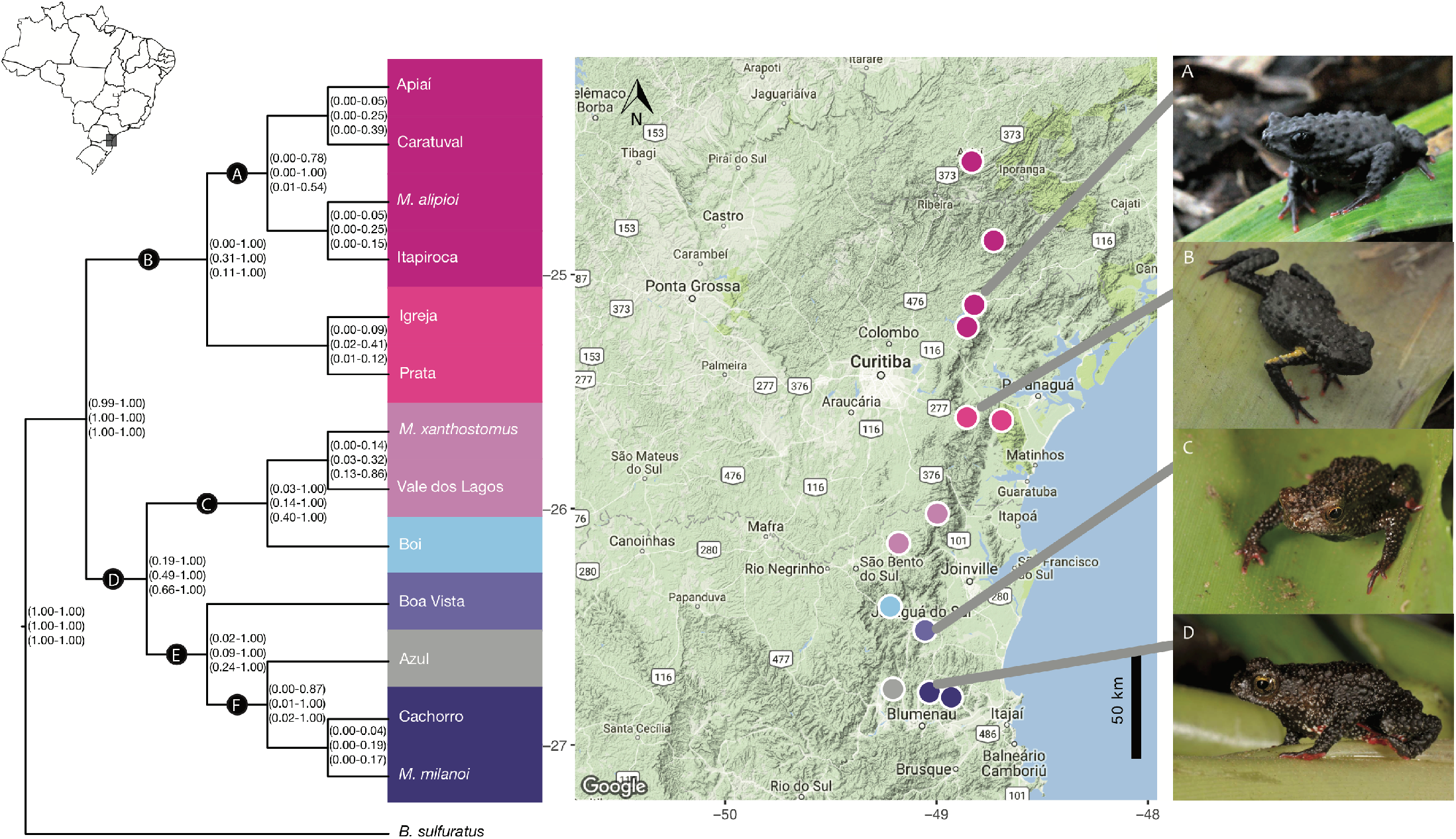
Species delimitation analyses of *Melanophryniscus* from the southern Brazilian Atlantic Forest. The three sets of values in parentheses indicate the range of posterior probabilities across all analyses for the corresponding node for each of three sets of priors [θ ~ Γ(2, 1000), τ0~ Γ(2, 1000); θ ~ Γ(2, 1000), τ0~ Γ(1, 10); and θ ~ Γ(1, 10), τ0~ Γ(1, 10)]. Nodes with considerable variation in posterior probabilities among analyses are indicated in black circles with letters. Colors correspond to the best-supported delimitation scheme. Background image from Map data ©2017 Google. A)*M. alipioi;* B)*M*. sp. “Igreja”; C)*M*. sp. “Boa Vista”; and D) *M. milanoi*.

**Figure 3.**
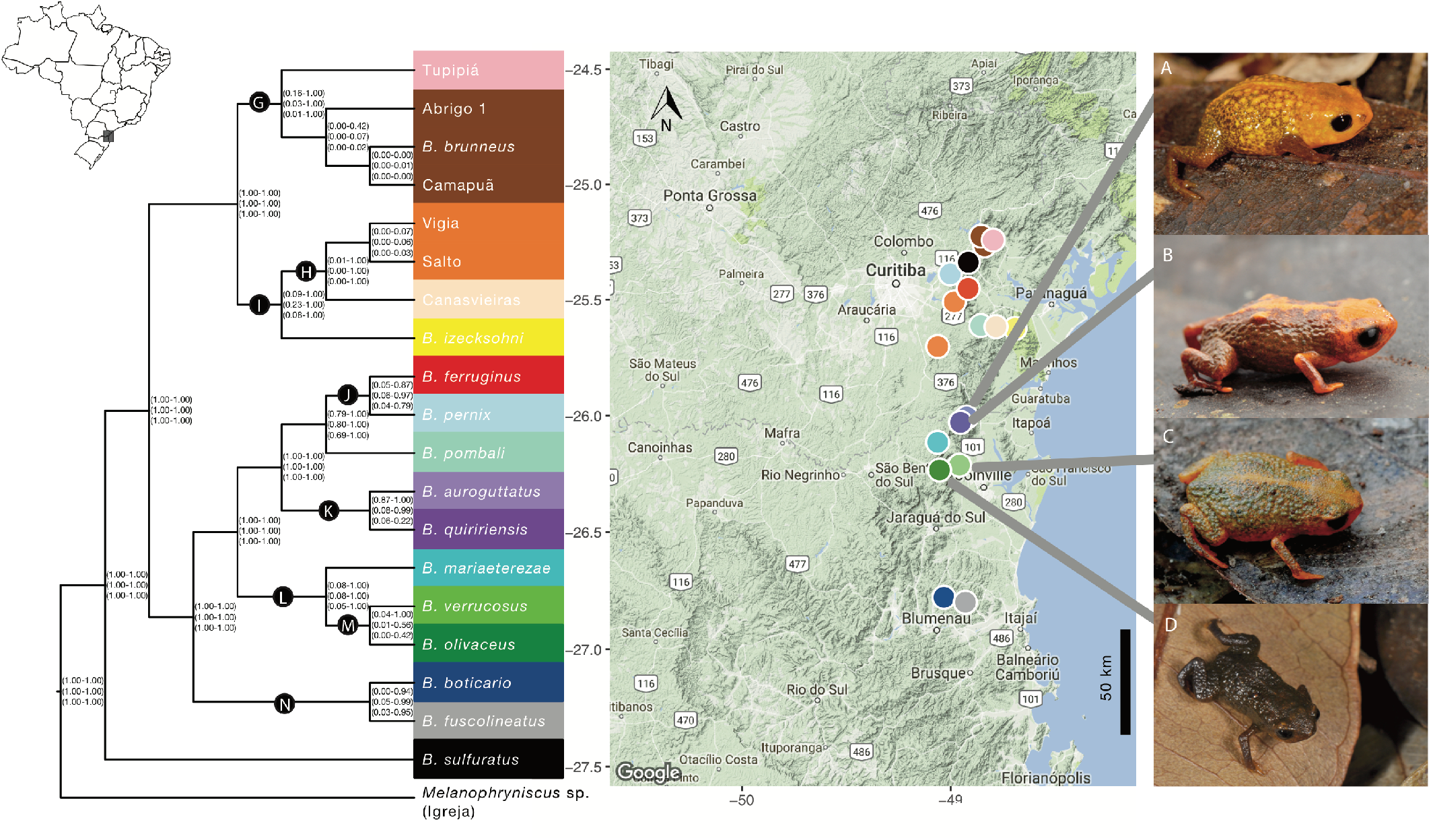
Species delimitation analyses of *Brachycephalus* from the southern Brazilian Atlantic Forest. The three sets of values in parentheses indicate the range of posterior probabilities across all analyses for the corresponding node for each of three sets of priors [θ ~ Γ(2, 1000), τ0~ Γ(2, 1000); θ ~ Γ(2, 1000), τ0~ Γ(1, 10); and θ ~ Γ(1, 10), τ0~ Γ(1, 10)]. Nodes with considerable variation in posterior probabilities among analyses are indicated in black circles with letters. Colors correspond to the best-supported delimitation scheme. Background image from Map data ©2017 Google. A) *B. auroguttatus;* B) *B. quiririensis;* C) *B. verrucosus;* and D) *B. olivaceus*.

**Figure 4.**
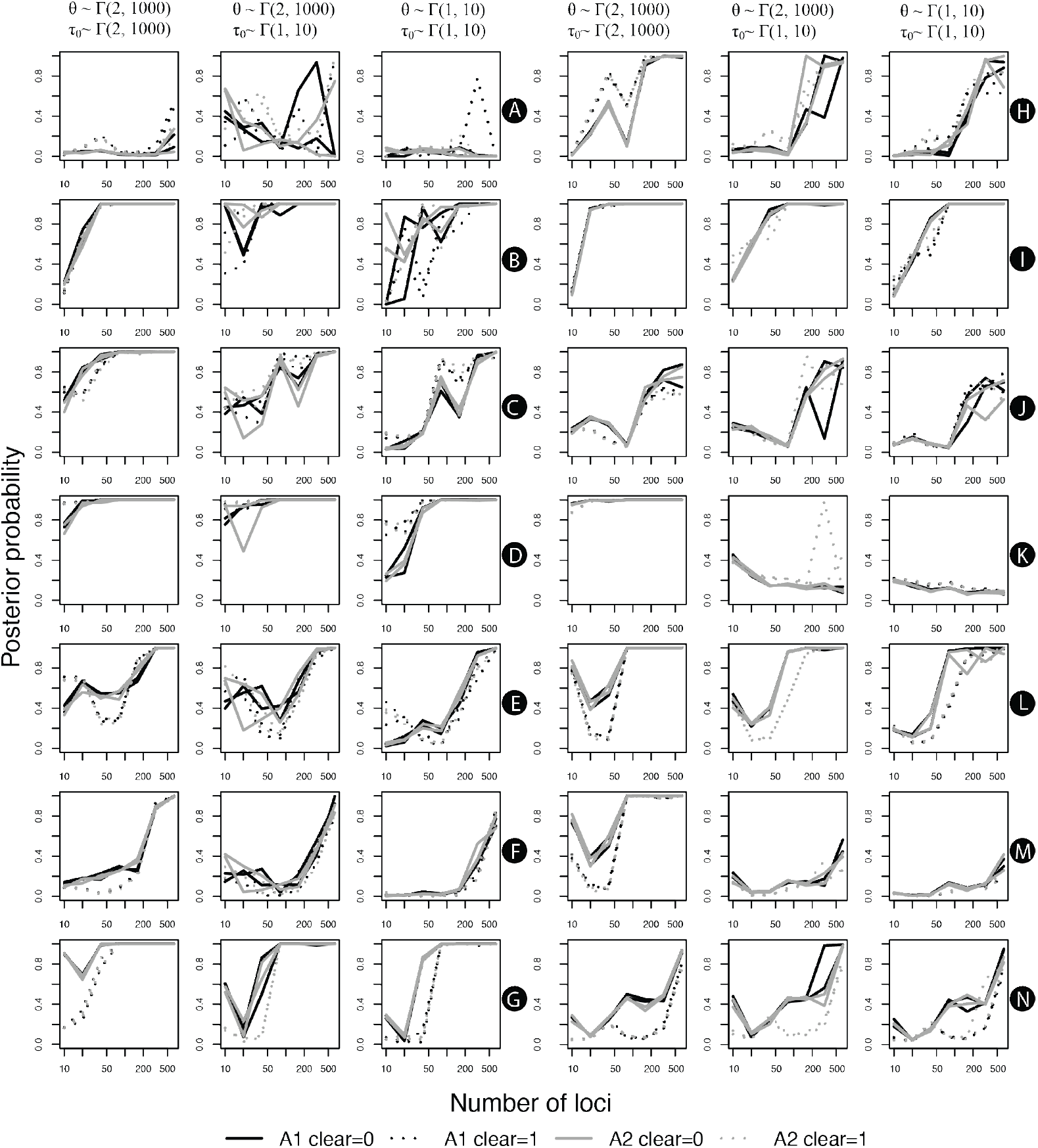
Variation in the posterior probability of lineage differentiation based on BPP species delimitation analyses according to the number of analyzed loci. Letters in black circles indicate the corresponding nodes on Figures 2 and 3. Line colors indicate results using either algorithm (black for algorithm 1, gray for algorithm 2), whereas line types indicate the treatment of sites with gaps (not omitted in solid lines, omitted in dotted lines). Different lines with the same color and type indicate different replicates of the same analysis.

A comparison of the results from ranked and randomly selected loci is shown in Figure 5. To facilitate their comparison, we present only the mean PPs between replicates for the ranked loci, and the mean PPs among the 10 randomly selected loci datasets. In general, the effect of adding more loci had a stronger impact on the variation in PPs than ranking them, given that the resulting PPs were consistently closer to the results of the 640-locus dataset. This effect seems to be mostly apparent up to the 160-loci datasets (Figure 5), but this threshold has to be interpreted with caution, given that randomly selected datasets tend to share increasingly more similar with the ranked dataset as more loci are added simply as a result of the finite total dataset.

**Figure 5.**
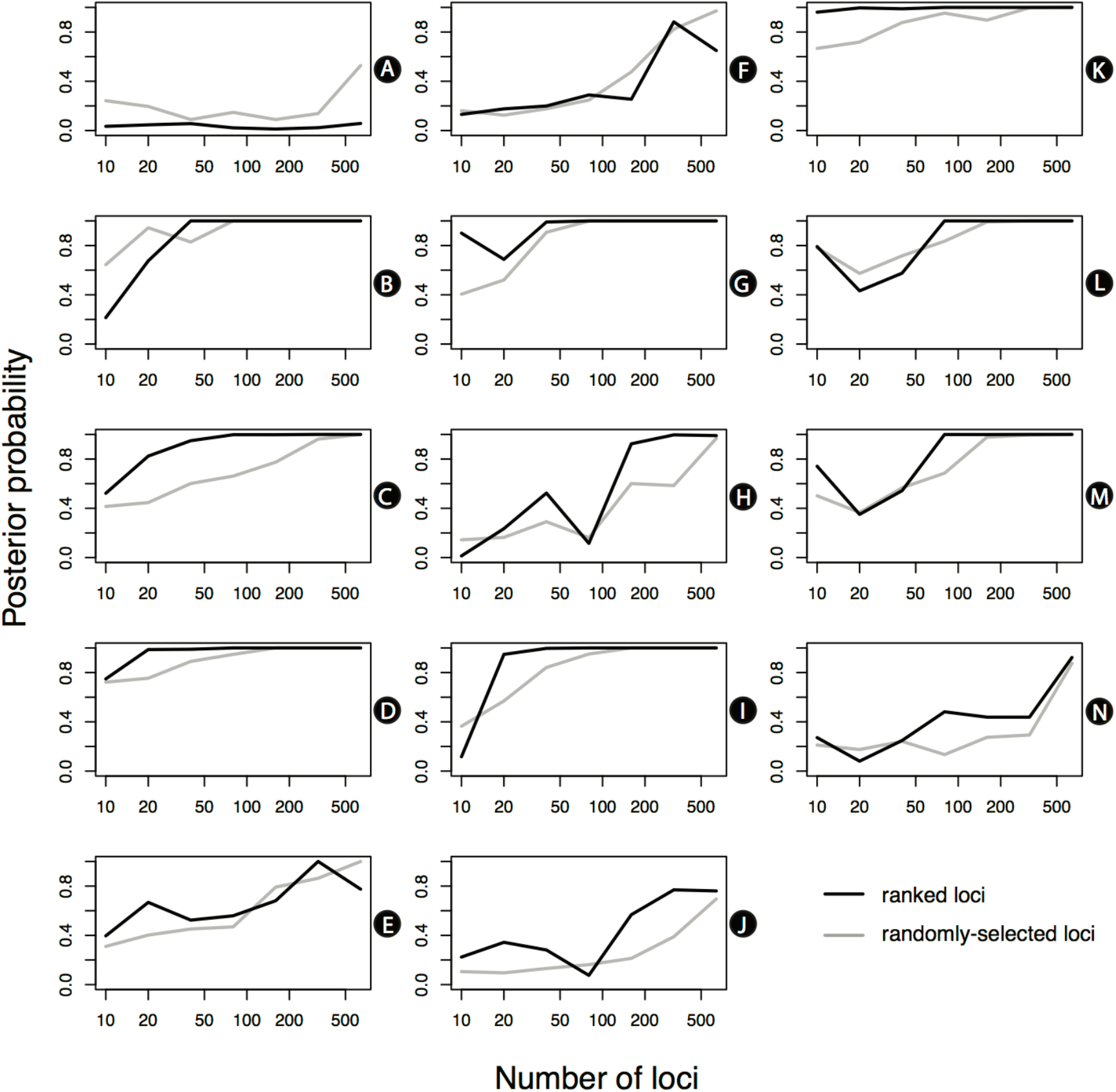
Variation in the posterior probability of lineage differentiation based on BPP species delimitation analyses according to the number of ranked and randomly selected loci. Letters in black circles indicate the corresponding nodes on Figures 2 and 3. To facilitate their comparison, we present only the mean PPs between replicates for the ranked loci, and the mean PPs among the 10 randomly selected loci datasets.

## Discussion

We carried out an extensive exploration of phylogenomic UCE data for species delimitation. Phylogenomic data are still under-utilized for species delimitation (Pyron 2015; but see Herrera and Shank 2016; de Oca et al. 2017). Indeed, to the best of our knowledge, only three other studies on molecular species delimitation have used ultraconserved elements (i.e. Smith et al. 2013; Newman and Austin 2016; Oswald et al. 2016). Newman and Austin (2016) used two alternative datasets: one with the 70 most informative loci, and a second dataset with 70 randomly selected loci from the full set of informative loci. In the first dataset, only one set of priors led to convergence (runs with small ancestral population size, i.e. θ = 2, 2000), whereas on the second dataset did not reach convergence, even after 300 k iterations. Oswald et al. (2016) used Bayes factor species delimitation using SNPs drawn from UCEs with the BFD* method (Grummer et al. 2014; Leaché et al. 2014). Given that only one SNP was used per locus to minimize genetic data linkage, the difference in their chosen method means that their results are not directly comparable to those in this study. Nevertheless, as UCEs and other types of genomic tool become increasingly more available (e.g. Starrett et al. in press; Branstetter et al. in press, Faircloth in press), we anticipate that phylogenomic data might become a valuable tool for species discovery, delimitation, and diagnosis. In particular, our results are consistent with Hime et al. (2016) in that, although species often can be delimited with relatively few loci, sampling larger numbers of loci might be necessary to ensure that recent divergences are correctly identified. In that study, Hime et al. explored variation from 2 to 10 loci using SpedeSTEM (Ence and Carstens 2011) and from 10 to 89 loci in analyses using BPP, in 10-locus increments. In those analyses, for each subsampling increment, they generated 10 replicate jackknifed datasets. They also ranked loci based on their informativeness, with increasing numbers of loci (as in the present study). Both the SpedeSTEM and BPP analyses showed more fine-scale delimitation with increasing number of loci, as expected based on simulation studies (e.g. Camargo et al. 2012; Hird et al. 2010). Indeed, our results strongly suggest that increasing the number of loci had a greater effect on delimitation than ranking them based on informativeness and focusing on the most informative loci, which suggests that uninformative loci might be more problematic for phylogenetic inference than for species delimitation analyses (at least in the case of BPP).

The use of genetic data for species delimitation has been recently criticized by Sukumaran and Knowles (2017), who argued that species delimitation based on the MSC, particularly as implemented in BPP, diagnoses genetic structure, not species, and that it cannot statistically distinguish structure associated with population isolation vs. species boundaries. The main concern raised by Sukumaran and Knowles (2017) is the possibility of taxonomic inflation if species are described based on genetic data alone (see also Olave et al. 2014). We believe that their concern is highly relevant in theory, but a relatively moot point in practice for two main reasons. First, although many studies in recent years have used genetic species delimitation methods, very few actual species descriptions have been based on genetic data alone. (Incidentally, in possibly the only instance to date of species descriptions based exclusively on BPP species delimitations, Leaché and Fujita [2010] have been mostly criticized by their lack of conformity with the International Code of Zoological Nomenclature [Bauer et al. 2010], although the possibility of taxonomic inflation is also raised.) Rather, nearly all studies using molecular species delimitation that led to actual species descriptions used other sources of data as well. This is perhaps surprising, given the consistent admonitions for the use of other types of data (morphological, behavioral, ecological) in conjunction with molecular data in species descriptions (e.g. Bauer et al. 2010; Rittmeyer and Austin 2012; Solís-Lemus et al. 2015).

Second, although conservation and management efforts ordinarily use species numbers as their main currency, such that taxonomic inflation could potentially lead to flawed conclusions, it is important to note that the opposite problem (the failure to recognize cryptic species as distinct entities) is at least equally as important (Bickford et al. 2007). Molecular species delimitation is often used precisely when diagnosis using phenotypic traits alone is difficult. It is therefore ironic that the recommendation is to go back to the phenotype, which is often (tacitly) assumed to be more reliable. Yet one might easily forget that phenotypic traits are often plagued with the same kinds of uncertainty as those indicated for molecular data, such as the difficulty in finding fixed diagnostic traits among potentially distinct species (Wiens and Servedio 2000), and the challenge of discriminating between differentiated populations from “good” species. It is possible that some of the criticism of molecular species delimitation methods has been due to the naive optimism of some early efforts that it would finally provide an objective means of defining species (as in the early days of molecular taxonomy), but the practice of taxonomic research has already demonstrated that species delimitation will never be an automated task, with or without phenotypic traits. In the end, it is important to keep in mind that cryptic species do exist, and that their biological reality should not be negated simply because fixed phenotypic diagnostic traits have not been identified.

Our results strongly corroborate the evolutionary distinctiveness of anuran species that had been previously described using phenotypic data alone (e.g. Bornschein et al. 2015, 2016b; Pie and Ribeiro 2015; Ribeiro et al. 2015). This is significant, given that many of those species — particularly in the case of *Brachycephalus* — have been diagnosed to a large extent based only on coloration, a trait that is often considered unreliable for anuran taxonomy. In addition, our analyses found evidence for 3 undescribed species of *Brachycephalus* and that three currently recognized *Melanophryniscus* species (i.e. *M. alipioi, M. xanthostomus*, and *M. milanoi*) actually represent species complexes involving a total of seven species. Interestingly, three pairs of closely related species (e.g. *B. ferruginus* X *B. pernix, B. auroguttatus* X *B. quiririensis*, and *B. verrucosus* X *B. olivaceus* – see Figure 3) did not reach 1.0 posterior probability in all analyses, despite having clear morphological diagnoses (including color variation), suggesting that BPP might in fact be conservative in its species delimitation compared to phenotypic evidence. The confirmation of the genomic distinctiveness of the montane anuran species in our study that had been previously described using phenotypic data alone, and the possibility of additional new species that require further study, has important implications for their conservation. Cloud forests, which are the habitat of most of the species in the *pernix* group, are among the most threatened ecosystems globally (Doumenge et al. 1995; Aldrich et al. 1997; Toledo-Aceves et al. 2011). This is of particular concern, not only due to the key role played by these forests in hydrological cycle maintenance, but also because they are reservoirs of endemic biodiversity (Toledo-Aceves et al. 2011). Many of the species in this study are categorized as either threatened or as data deficient, yet any level of formal governmental protection necessarily involves the availability of a species name. Given their microendemic distribution and highly threatened habitats, one could argue that commission errors (considering them as different species) would be preferable in relation to omission errors (lumping them as single species). In practice, urgent management efforts should be enforced to ensure their long-term survival.

## Acknowledgements

This study was partially funded by a grant from Fundação Grupo Boticário de Proteção à Natureza (grant # 0895_20111). Samples were collected under ICMBIO permit #22470-2 and Instituto Ambiental do Paraná permit 355/11. MRP was funded by CNPq/MCT (grant 571334/2008-3). Whitney Tsai provided invaluable assistance for obtaining UCE data.

## Supplementary material

**Figure S1.**
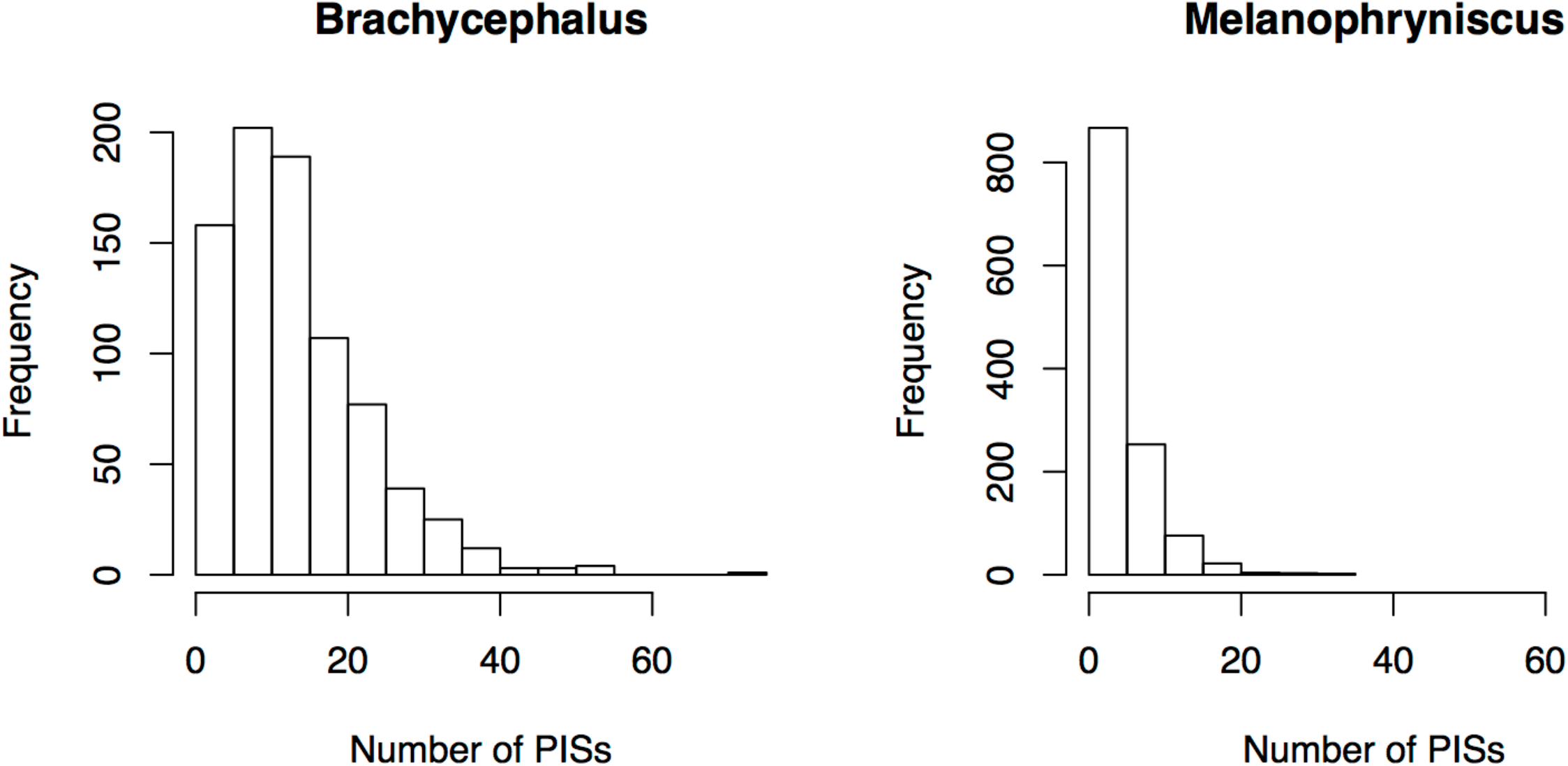
Frequency distribution of the number of parsimony-informative sites (PISs) in the *Brachycephalus* and *Melanophryniscus* datasets (N=820 and 1227 loci, respectively).

## References

Adams M., Raadik T.A., Burridge C.P., Georges A. 2014. Global biodiversity assessment and hyper-cryptic species complexes: more than one species of elephant in the room? Syst. Biol. 63:518–533.

Aldrich M., Billington C., Edwards M., Laidlaw R. 1997. Tropical montane cloud forests: an urgent priority for conservation. WCMC Biodiversity Bulletin 2:1–14.

Bauer A.M., Parham J.F., Brown R.M., Stuart B.L., Grismer L., Papenfuss T.J., Böhme W., Savage, C.S., Grismer J.L., Wagner, P., Schmitz A., Ananjeva N.B., Inger, R.F., Wagner, P. 2010. Availability of new Bayesian-delimited gecko names and the importance of character-based species descriptions. Proc. R. Soc. B 278:490–492.

Bi K., Vanderpool D., Singhal S., Linderoth, T., Moritz, C., Good, J.M. 2012. Transcriptome-based exon capture enables highly cost-effective comparative genomic data collection at moderate evolutionary scales. BMC Genomics 13: 403.

Bickford D., Lohman D.J., Sodhi N.S., Ng P.K., Meier R., Winker K., Ingram, K.K., Das, I. 2007. Cryptic species as a window on diversity and conservation. Trends Ecol. Evol. 22:148–155.

Blumenstiel B., Cibulskis K., Fisher S., DeFelice M., Barry A., Fennell T., Abreu J., Minie B., Costello M., Young G., Maguire J., Melnikov A., Rogov P., Gnirke A., Gabriel S. 2010. Targeted exon sequencing by in-solution hybrid selection. Curr. Protoc. Hum. Genet. Chapter 18. Unit 18.4.

Bolger A.M., Lohse M., Usadel B. 2014. Trimmomatic: a flexible trimmer for Illumina sequence data. Bioinformatics 30:2114–2120

Bornschein M.R., Firkowski C.R., Baldo D., Ribeiro L.F., Belmonte-Lopes R., Corrêa L., Morato S.A.A., Pie M.R. 2015. Three new species of phytotelm-breeding *Melanophryniscus* from the Atlantic Rainforest of southern Brazil (Anura: Bufonidae). Plos One 10:e0142791.

Bornschein M.R., Firkowski C.R., Belmonte-Lopes R., Corrêa L., Ribeiro L.F., Morato S.A., Antoniazzi-Jr R.L., Reinert B.L., Meyer A.L., Cini F.A., Pie M.R. 2016a. Geographical and altitudinal distribution of *Brachycephalus* (Anura: Brachycephalidae) endemic to the Brazilian Atlantic Rainforest. PeerJ 4:e2490.

Bornschein M.R., Ribeiro L.F., Blackburn D.C., Stanley E.L., Pie M.R. 2016b. A new species of *Brachycephalus* (Anura: Brachycephalidae) from Santa Catarina, southern Brazil. PeerJ. 4:e2629.

Branstetter M.G., Longino J.T., Ward P.S., Faircloth, B.C. in press. Enriching the ant tree of life: enhanced UCE bait set for genome-scale phylogenetics of ants and other Hymenoptera. Methods Ecol. Evol.

Camargo A., Morando M., Avila L.J., Sites J.W. Jr. 2012. Species delimitation with ABC and other coalescent-based methods: a test of accuracy with simulations and an empirical example with lizards of the *Liolaemus darwinii* complex (Squamata:Liolaemidae). Evolution 66:2834–2849.

Castresana J. 2000. Selection of conserved blocks for multiple alignments for their use in phylogenetic alignments. Mol. Biol. Evol. 17:540–552.

Caviedes-Solis I.W., Bouzid N.M., Banbury B.L., Leaché A.D. 2015. Uprooting phylogenetic uncertainty in coalescent species delimitation: A meta-analysis of empirical studies. Cur. Zool. 61:866–73.

de Oca A.N.M., Barley A.J., Meza-Lázaro R.N., García-Vázquez U.O., Zamora-Abrego J.G., Thomson R.C., Leaché A.D. 2017. Phylogenomics and species delimitation in the knob-scaled lizards of the genus *Xenosaurus* (Squamata: Xenosauridae) using ddRADseq data reveal a substantial underestimation of diversity. Mol. Phylogenet. Evol. 106:241–253.

Domingos F.M., Colli G.R., Lemmon A., Lemmon E.M., Beheregaray L.B. 2017. In the shadows: Phylogenomics and coalescent species delimitation unveil cryptic diversity in a Cerrado endemic lizard (Squamata: *Tropidurus*). Mol. Phylogenet. Evol. 107:455–465.

Doumenge C., Gilmour D., Pérez, M.R., Blockhus, J. 1995. Tropical montane cloud forests: conservation status and management issues. In Tropical montane cloud forests (pp. 24–37). Springer US.

Ence D.D., Carstens B.C. 2011. spedeSTEM: a rapid and accurate method for species delimitation. Mol. Ecol. Res. 11:473–480.

Faircloth B.C. 2015. PHYLUCE is a software package for the analysis of conserved genomic loci. Bioinformatics 32:786–788.

Faircloth B.C. in press. Identifying conserved genomic elements and designing universal bait sets to enrich them. Methods Ecol. Evol.

Faircloth B.C., Glenn T.C. 2012. Not all sequence tags are created equal: designing and validating sequence identification tags robust to indels. PLoS One 7:e42543.

Faircloth, B.C., McCormack, J.E., Crawford, N.G., Harvey, M.G., Brumfield, R.T., Glenn, T.C. 2012. Ultraconserved elements anchor thousands of genetic markers spanning multiple evolutionary timescales. Syst. Biol. 61:717–726.

Firkowski C.R., Bornschein M.R., Ribeiro L.F., Pie M.R. (2016). Species delimitation, phylogeny and evolutionary demography of co-distributed, montane frogs in the southern Brazilian Atlantic Forest. Mol. Phylogenet. Evol. 100:345–360.

Frost, D.R. 2017. Amphibian species of the world: an online reference. Electronic Database (accessed January, 2017).

Fujita, M.K., Leaché, A.D. 2011. A coalescent perspective on delimiting and naming species: a reply to Bauer et al. Proc. R. Soc. Lond. B Biol. Sci. 278:493–495.

Gilbert P.S., Chang J., Pan C., Sobel E.M., Sinsheimer J.S., Faircloth B.C., Alfaro M.E. 2015. Genome-wide ultraconserved elements exhibit higher phylogenetic informativeness than traditional gene markers in percomorph fishes. Mol. Phylogenet. Evol. 92:140–146.

Gnirke A., Melnikov A., Maguire J., Rogov P., LeProust E.M., Brockman W., Fennell T., Giannoukos G., Fisher S., Russ C., Gabriel S., Jaffe D.B., Lander E.S., Nusbaum C. 2009. Solution hybrid selection with ultra-long oligonucleotides for massively parallel targeted sequencing. Nat. Biotechnol. 27:182–189.

Grabherr, M.G., Haas, B.J., Yassour, M. et al. 2011. Full-length transcriptome assembly from RNA-Seq data without a reference genome. Nature Biotechnol. 29:644–652.

Grummer J.A., Bryson R.W. Reeder T.W. 2014. Species delimitation using Bayes factors: simulations and application to the *Sceloporus scalaris* species group (Squamata: Phrynosomatidae). Syst. Biol. 63:119–133.

Hanken J., Wake, D.B. 1993. Miniaturization of body size: organismal consequences and evolutionary significance. Annu. Rev. Ecol. Syst. 24:501–519.

Heibl C. 2014. ips: Interfaces to Phylogenetic Software in R. Available from CRAN at https://CRAN.R-project.org/package=ips

Herrera S., Shank T.M. 2016. RAD sequencing enables unprecedented phylogenetic resolution and objective species delimitation in recalcitrant divergent taxa. Mol. Phylogenet. Evol. 100:70–79.

Hime P.M., Hotaling S., Grewelle R.E., O’Neill E.M., Voss S.R., Shaffer H.B., Weisrock D.W. 2016. The influence of locus number and information content on species delimitation: an empirical test case in an endangered Mexican salamander. Mol. Ecol. 25:5959–5974.

Hird S., Kubatko L., Carstens, B. 2010. Rapid and accurate species tree estimation for phylogeographic investigations using replicated subsampling. Mol. Phylogenet. Evol. 57:888–898.

Hosner P.A., Faircloth B.C., Glenn T.C., Braun E.L., Kimball R.T. 2016. Avoiding missing data biases in phylogenomic inference: an empirical study in the landfowl (Aves: Galliformes). Mol. Biol. Evol. 33:1110–1125.

Jarvis E.D., Mirarab S., Aberer A.J. et al. 2014 Whole-genome analyses resolve early branches in the tree of life of modern birds. Science 346:1320–1331.

Katoh S. 2013. MAFFT multiple sequence alignment software version 7: improvements in performance and usability. Mol. Biol. Evol. 30:772–780

Knowles L.L., Carstens B.C. 2007. Delimiting species without monophyletic gene trees. Syst. Biol. 56:887–895.

Langone J.A., Segalla M.V., Bornschein M.R., de Sá R.O. 2008. A new reproductive mode in the genus *Melanophryniscus* Gallardo, 1961 (Anura: Bufonidae) with description of a new species from the state of Paraná, Brazil. S. Am. J. Herpetol. 3:1–9.

Leaché A.D., Fujita M.K. 2010. Bayesian species delimitation in West African forest geckos (*Hemidactylus fasciatus)*. Proc. R. Soc. Lond. B Biol. Sci. 277:3071–3077.

Leaché A.D., Fujita M.K., Minin, V.N. Bouckaert R.R. 2014. Species delimitation using genome-wide SNP data. Syst. Biol. 63:534–542.

Lemmon A.R., Emme S.A., Lemmon E.M. 2012. Anchored hybrid enrichment for massively high-throughput phylogenomics. Syst. Biol. 61:727–44.

Lemmon E.M., Lemmon A.R. 2013. High-throughput genomic data in systematics and phylogenetics. Annu. Rev. Ecol. Evol. Syst. 44:99–121.

Manthey J.D., Campillo L.C., Burns K.J., Moyle, R. G. 2016. Comparison of target-capture and restriction-site associated DNA sequencing for phylogenomics: a test in cardinalid tanagers (Aves, Genus: *Piranga*). Syst. Biol. 65:640–650.

McCormack J. E., Hird S.M., Zellmer A.J., Carstens B.C., Brumfield R.T. (2013). Applications of next-generation sequencing to phylogeography and phylogenetics. Mol. Phylogenet. Evol. 66:526–538.

Meiklejohn K.A., Faircloth B.C., Glenn T.C., Kimball R.T., Braun E.L. 2016. Analysis of a rapid evolutionary radiation using ultraconserved elements: evidence for a bias in some multispecies coalescent methods. Syst. Biol. 65:612–627

Newman, C.E., Austin, C.C. 2016. Sequence capture and next-generation sequencing of ultraconserved elements in a large-genome salamander. Mol. Ecol. 25:6162–6174.

Olave M., Solà E., Knowles L.L., 2014. Upstream analyses create problems with DNA-based species delimitation. Syst. Biol 63:263–271.

Oswald J.A., Harvey M.G., Remsen R.C., Foxworth D.U., Cardiff S.W., Dittmann D.L., Megna L.C., Carling M.D., Brumfield R.T. 2016. Willet be one species or two? A genomic view of the evolutionary history of *Tringa semipalmata*. The Auk 133:593–614.

Pie M.R., Meyer A.L., Firkowski C.R., Ribeiro L.F., Bornschein M.R. 2013. Understanding the mechanisms underlying the distribution of microendemic montane frogs (*Brachycephalus* spp., Terrarana: Brachycephalidae) in the Brazilian Atlantic Rainforest. Ecol. Model. 250:165–176.

Pie M.R. and Ribeiro L.F. 2015. A new species of *Brachycephalus* (Anura: Brachycephalidae) from the Quiriri mountain range of southern Brazil. PeerJ 3:e1179.

Pyron R.A. 2015. Post-molecular systematics and the future of phylogenetics. Trends Ecol. Evol. 30:384–389.

R Core Team. 2017. R: A language and environment for statistical computing. R Foundation for Statistical Computing, Vienna, Austria. URL https://www.R-project.org/.

Rannala B. 2015. The art and science of species delimitation. Curr. Zool. 61:846–853.

Rannala B., Yang Z. 2003. Bayes estimation of species divergence times and ancestral population sizes using DNA sequences from multiple loci. Genetics 164:1645–1656.

Rannala B., Yang Z. 2013. Improved reversible jump algorithms for Bayesian species delimitation. Genetics 194:245–253.

Ribeiro L.F., Bornschein M.R., Belmonte-Lopes R., Firkowski C.R., Morato S.A.A., Pie M.R. 2015. Seven new microendemic species of *Brachycephalus* (Anura: Brachycephalidae) from southern Brazil. PeerJ 3:e1011

Rittmeyer E.N., Austin C.C. 2012. The effects of sampling on delimiting species from multi-locus sequence data. Mol. Phylogenet. Evol 65:451–463.

Sites Jr, J.W., Marshall J.C. 2004. Operational criteria for delimiting species. Annu. Rev. Ecol. Evol. Syst. 35:199–227.

Smith B.T., Harvey M.G., Faircloth B.C., Glenn T.C., Brumfield R.T. 2013. Target capture and massively parallel sequencing of ultraconserved elements (UCEs) for comparative studies at shallow evolutionary time scales. Syst. Biol. 63:83–95.

Solís-Lemus C., Knowles L.L., Ané C. 2015. Bayesian species delimitation combining multiple genes and traits in a unified framework. Evolution 69:492–507.

Starrett J., Derkarabetian S., Hedin M., Bryson, R.W., McCormack J.E., Faircloth B.C. in press. High phylogenetic utility of an ultraconserved element probe set designed for Arachnida. Mol. Ecol. Resour.

Steinbach-Padilha G.C. 2008. A new species of *Melanophryniscus* (Anura, Bufonidae) from the Campos Gerais region of Southern Brazil. Phyllomedusa 7:99–108.

Sukumaran J., Knowles L.L. 2017. Multispecies coalescent delimits structure, not species. Proc. Natl. Acad. Sci. USA 114:1607–1612.

Takahata N., Satta Y., Klein J. 1995. Divergence time and population-size in the lineage leading to modern humans. Theor. Pop. Biol. 48:198–221.

Toledo-Aceves T., Meave J.A., González-Espinosa M., Ramírez-Marcial N. 2011. Tropical montane cloud forests: current threats and opportunities for their conservation and sustainable management in Mexico. J. Environ. Manageme. 92:974–981.

Townsend J. P. 2007). Profiling phylogenetic informativeness. Syst. Biol, 56:222–231.

Wiens J.J., Servedio M.R. 2000. Species delimitation in systematics: inferring diagnostic differences between species. Proc. R. Soc. Lond. B Biol. Sci. 267:631–636.

Xi Z., Liu L., Davis C.C. 2015. Genes with minimal phylogenetic information are problematic for coalescent analyses when gene tree estimation is biased. Mol. Phylogenet. Evol. 92:63–71.

Xu B., Yang Z. 2016. Challenges in species tree estimation under the multispecies coalescent model. Genetics 204:1353–1368.

Yang Z. 2002. Likelihood and Bayes estimation of ancestral population sizes in Hominoids using data from multiple loci. Genetics 162:1811–1823.

Yang Z. 2015. The BPP program for species tree estimation and species delimitation. Curr. Zool. 61:854–865.

Yang Z., Rannala, B. 2010. Bayesian species delimitation using multilocus sequence data. Proc. Natl. Acad. Sci. U.S.A. 107:9264–9269.

Yang Z., Rannala B. 2014. Unguided species delimitation using DNA sequence data from multiple loci. Mol. Biol. Evol. 31:3125–3135.

Yeh J. 2002. The effect of miniaturized body size on skeletal morphology in frogs. Evolution 56:628–641.

Zhang C., Rannala B., Yang Z. 2014. Bayesian species delimitation can be robust to guide-tree inference errors. Syst. Biol. 63:993–1004.

Zhang C., Zhang D.X., Zhu T., Yang Z. 2011. Evaluation of a Bayesian coalescent method of species delimitation. Syst. Biol. 60:747–761.

